# Identification of different putative outer membrane electron conduits necessary for Fe(III) citrate, Fe(III) oxide, Mn(IV) oxide, or electrode reduction by *Geobacter sulfurreducens*

**DOI:** 10.1101/169086

**Authors:** Fernanda Jiménez Otero, Chi Ho Chan, Daniel R. Bond

## Abstract

At least five gene clusters in the *Geobacter sulfurreducens* genome encode putative ‘electron conduits’ implicated in electron transfer across the outer membrane, each containing a periplasmic multiheme *c*-type cytochrome, integral outer membrane anchor, and outer membrane redox lipoprotein(s). Markerless single gene cluster deletions and all possible multiple deletion combinations were constructed and grown with soluble Fe(III) citrate, Fe(III)- and Mn(IV)-oxides, and graphite electrodes poised at +0.24 V and −0.1 V vs. SHE. Different gene clusters were necessary for reduction of each electron acceptor. During metal oxide reduction, deletion of the previously described *omcBC* cluster caused defects, but deletion of additional components in an Δ*omcBC* background, such as *extEFG*, were needed to produce defects greater than 50% compared to wild type. Deletion of all five gene clusters abolished all metal reduction. During electrode reduction, only the Δ*extABCD* mutant had a severe growth defect at both redox potentials, while this mutation did not affect Fe(III)-oxide, Mn(IV)-oxide, or Fe(III) citrate reduction. Some mutants containing only one cluster were able to reduce particular terminal electron acceptors better than wild type, suggesting routes for improvement by targeting specific electron transfer pathways. Transcriptomic comparisons between fumarate and electrode-based growth showed all of these *ext* clusters to be constitutive, and transcriptional analysis of the triple-deletion strain containing only *extABCD* detected no significant changes in expression of known redox proteins or pili components. These genetic experiments reveal new outer membrane conduit complexes necessary for growth of *G. sulfurreducens*, depending on the available extracellular electron acceptor.

## IMPORTANCE

Gram-negative metal-reducing bacteria utilize electron conduits, chains of redox proteins spanning the outer membrane, to transfer electrons to the extracellular surface. Only one pathway for electron transfer across the outer membrane of *Geobacter sulfurreducens* has been linked to Fe(III) reduction. However, *G. sulfurreducens* is able to respire a wide array of extracellular substrates. Here, we present the first combinatorial genetic analysis of five different electron conduits via creation of new markerless deletion strains and complementation vectors. Multiple conduit gene clusters appear to have overlapping roles, including two that have never been linked to metal reduction. Another recently described cluster (ExtABCD) was the only electron conduit essential during electrode reduction, a substrate of special importance to biotechnological applications of this organism.

## INTRODUCTION

Microorganisms capable of extracellular respiration can alter the redox state of particulate metal oxides in soils and sediments, controlling their solubility and bioavailability (1–6). To respire with extracellular metals, bacteria must first transfer electrons from the cell interior to outer surface redox proteins, requiring unique transmembrane pathways compared to growth with intracellularly-reduced compounds. The use of surface-exposed electron transfer proteins and conductive appendages by these organisms presents opportunities for transformation of heavy metals, biological nanoparticle synthesis, and a new generation of microbially-powered electrochemical devices using bacteria grown on electrodes (7–13).

An extracellular electron transfer strategy must overcome several challenges. In Gram-negative cells, a conductive pathway capable of crossing the inner membrane, periplasm, and outer membrane must first be constructed (14, 15). Such pathways are capable of delivering electrons to soluble metals or redox-active molecules, but insoluble metal oxides present additional barriers. Fe(III)- and Mn(IV)-oxides vary widely in chemistry, surface charge, redox state, and surface area, thus an additional suite of proteins or appendages such as pili may be needed to link cell surfaces with different terminal minerals (16–18).

Many metal-reducing bacteria can also transfer electrons to electrodes (8, 19–21). Unlike metal oxide particles, electrodes represent an unlimited electron acceptor allowing cells in contact with the inorganic surface to support growth of more distant cells, if they can create a conductive network of proteins that relay electrons to cells at the electrode. The physiological and chemical differences between soluble metals, metal particles, and electrodes raises the possibility that different electron transfer proteins may be needed to access each kind of extracellular mineral, surface, or molecule.

A model organism widely studied for its ability to reduce a diversity of metals and electrodes is the δ-Proteobacterium *Geobacter sulfurreducens*, and recent work supports a model where different electron transfer proteins are used depending on substrate conditions. At the inner membrane where electrons first leave the quinone pool, a combination *c-* and *b*-type cytochrome CbcL (22) is only required when extracellular metals and electrodes are below redox potentials of −0.1 V vs. SHE, while the inner membrane *c*-type cytochrome ImcH (23), becomes essential if acceptors are at higher redox potentials (18). In another example, in the extracellular matrix beyond the cell surface, chemistry rather than redox potential appears to delineate which proteins are essential for electron transfer. The secreted cytochrome OmcZ and pili-based appendages are primarily linked to electrode growth, while the secreted cytochrome PgcA enhances reduction of Fe(III)-oxides without affecting electrode growth (24–31). Between the initial CbcL/ImcH-dependent event of inner membrane proton motive force generation and extracellular pili/OmcZ/PgcA interactions lies the outer membrane, a less understood barrier that was recently found to contain electron transfer proteins of surprising complexity (32–34).

The only known mechanism for non-diffusive electron transfer across the outer membrane is through a transmembrane ‘electron conduit’, consisting of an integral outer membrane protein anchoring a periplasmic multiheme cytochrome to an outer surface lipoprotein cytochrome. By linking redox active cofactors within a membrane-spanning complex, electron flow is permitted (32, 35). The first electron conduit described was the ~210 kDa MtrCAB complex from *S. oneidensis*, which will catalyze electron transfer across membranes when purified and placed in lipid vesicles (36–38). The *mtrCAB* gene cluster is essential for reduction of all tested soluble metals, electron shuttles, metal oxides, and electrodes by *S. oneidensis* (37, 39, 40). Related complexes capped with an extracellular DMSO reductase allow *Shewanella* to reduce DMSO on the cell exterior, while similar outer membrane conduits support inward electron flow by Fe(II)-oxidizing *Rhodopseudomonas* TIE-1 cells (41, 42).

In *G. sulfurreducens*, a gene cluster encoding the periplasmic cytochrome OmbB, the outer membrane integral protein OmaB, and lipoprotein cytochrome OmcB forms a conduit complex functionally similar to MtrCAB, though the two complexes lack any sequence similarity (34). This *‘ombB-omaB-omcB’* gene cluster is duplicated immediately downstream in the *G. sulfurreducens* genome as the near-identical *‘ombC-omaC-omcC’*, together forming the *‘omcBC* cluster. Antibiotic cassette insertions replacing *omcB*, as well as insertions replacing the entire *‘ombB-omaB-omcB’* conduit, decrease growth with Fe(III) as an electron acceptor, but the impact differs between reports and growth conditions (43–45). This variability and residual electron transfer activity suggested the presence of alternative pathways able to catalyze electron transfer across the outer membrane (33).

New evidence for undiscovered outer membrane complexes was recently detected in genome-wide transposon data, where insertions in *omcB* or *omcC* had no effect on *G. sulfurreducens* growth with electrodes poised at −0.1 vs. SHE, a low potential chosen to mimic the redox potential of Fe(III)-oxides (46). Transposon insertions within an unstudied four-gene cluster containing *c*-type cytochrome conduit signatures caused significant defects during growth on the same −0.1 V electrodes (46). Deletion of this new cluster, named *extABCD*, severely affected growth on low-potential electrodes, while *ΔextABCD* mutants still grew similar to wild type with Fe(III)-oxides. In contrast, deletion of the entire *omcBC* cluster had little impact on low-potential electrode growth (46). These data suggested that the outer membrane proteins essential for electron transfer across the membrane might vary depending on environmental conditions. However, these data involved only single deletions without complementation and did not test if different gene clusters were necessary across the full range of environmentally relevant conditions such as higher redox potentials, during growth with mineral forms such as Mn(VI), or when metals become soluble.

Using new markerless deletion methods, this study constructed mutants containing all combinations of the four putative conduit clusters on the genome of *G. sulfurreducens.* Each of these 15 mutants plus three strains containing expression vectors were then directly compared with five electron acceptors; Fe(III)- and Mn(IV)-oxides, poised electrodes at two different redox potentials, and soluble Fe(III)-citrate. We found that during metal reduction the largest defects were in Δ*omcBC* strains, but deletion of the newly identified cluster *extEFG* in the Δ*omcBC* background was necessary to most severely inhibit Fe(III)-reduction, and deletion of all clusters was required to eliminate reduction of both soluble and insoluble metals. Strains containing only a single cluster showed preferences for reduction of different metals, such as the *extEFG-* and *extHIJKL*-only strains performing better with Mn(IV)-oxides than Fe(III)-oxides. When electrodes were the electron acceptor, only strains lacking *extABCD* showed a growth defect, and this effect was similar at all redox potentials. A strain still containing *extABCD* but lacking all other conduit clusters grew faster and to a higher final density on electrodes, and a complemented strain lacking all other conduit clusters expressing *extABCD* from a vector also grew faster than wild type. These data provide evidence that different *G. sulfurreducens* conduit clusters are necessary during extracellular electron transfer depending on the extracellular substrate.

(This article was submitted to an online preprint archive (47))

## RESULTS

**Description of putative outer membrane electron conduit gene clusters**. At least five loci can be identified in the *G. sulfurreducens* genome encoding putative *c*-type cytochrome electron conduits. This identification is based on three key elements; (1) a multiheme periplasmic *c*-type cytochrome, (2) an outer membrane integral protein with transmembrane ß-sheets, and (3) one or more outer membrane lipoproteins with redox cofactors (Fig. 1A). Two of these regions correspond to the well-studied OmcB-based *(ombB-omaB-omcB*, GSU2739-2737) conduit and its near-identical duplicate OmcC-based operon immediately downstream preceded by a TetR-family repressor partially truncated in its DNA-binding domain *(orfS-ombC-omaC-omcC*, GSU2733-2731). For clarity, and due to the fact that *omaBC* and *ombBC* are identical, this region is referred to as the *“omcBC”* cluster. The well-characterized duplicate *omcBC* cluster was deleted as a single unit (see also Materials and Methods for additional information about the tendency of identical genes within this region to recombine during mutant construction and efforts taken to verify proper removal and reconstruction of *omcBC* genes).

**Figure 1.**
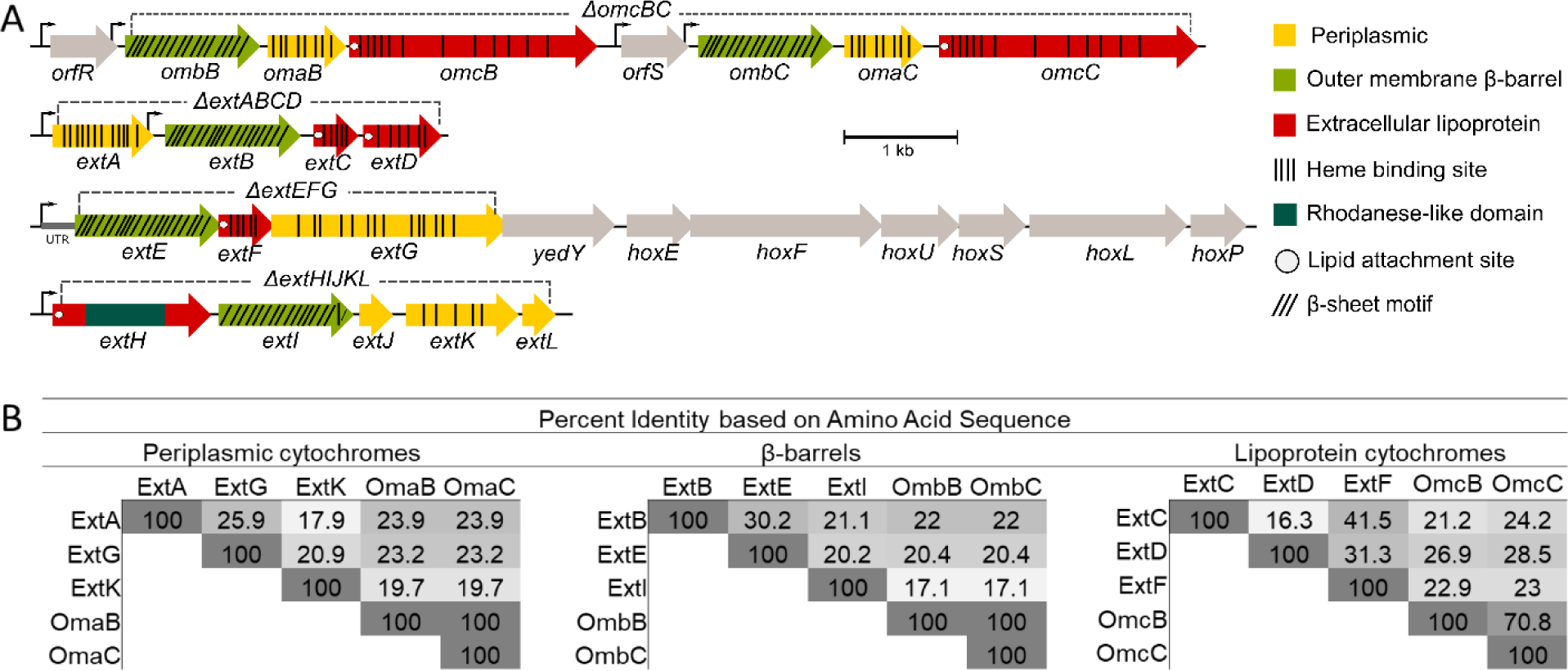
The outer membrane electron conduit gene clusters of *G. sulfurreducens*. A) Genetic organization and predicted features of operons containing putative outer membrane conduits. Deletion constructs indicated by dashed line. B) Identity matrix from amino acid sequence alignment of each cytochrome or β-barrel component using ClustalΩ.

The *ext* genes comprise three new clusters, named for their putative roles in <u>ext</u>racellular electron transfer (46). Relative protein orientations were predicted using a combination of protein localization prediction software (48), integral membrane prediction software (49), and lipid attachment site prediction software (50). The *extABCD* (GSU2645-2642) cluster encodes ExtA, a periplasmic dodecaheme *c*-type cytochrome, ExtB, an outer membrane integral protein with 18 trans-membrane domains, and ExtCD, two outer membrane lipoprotein *c*-type cytochromes with 5 and 12 heme binding sites, respectively. The second cluster, *extEFG* (GSU2726-2724), encodes ExtE, an outer membrane integral protein with 21 trans-membrane domains, ExtF, an outer membrane lipoprotein pentaheme *c*-type cytochrome, and ExtG, a periplasmic dodecaheme *c*-type cytochrome. Kanamycin insertions in ExtG have been shown to affect Fe(III)-oxide reduction, which in some annotations is referred to as OmcV (51) despite its predicted periplasmic localization. For consistency with the surrounding operon and to distinguish it from outer membrane cytochromes, the name ExtG will be used in this work. The final cluster, *extHIJKL* (GSU2940-2936) lacks an outer membrane *c*-type cytochrome, but encodes ExtH, a rhodanese-family lipoprotein, ExtI, a 21 trans-membrane domain outer membrane integral protein, ExtJ, a small periplasmic protein, and ExtKL, a periplasmic pentaheme *c*-type cytochrome followed by a small hypothetical protein. A TGA stop codon encoding a predicted rare selenocystine amino acid separates ExtK and ExtL, thus they may encode a single protein (52).

A significant difference between *G. sulfurreducens* Ext clusters and the *S. oneidensis* Mtr conduits (35), is that the *mtr* clusters in *S. oneidensis* are paralogs. The periplasmic MtrA and MtrD cytochromes share over 50% identity, are similar in size and heme content, and can cross-complement (53). The lipoprotein outer surface cytochromes of *Shewanella* also demonstrate high sequence, functional, and structural conservation (32, 53–55). In contrast, no component of the Ext or OmcB complexes share any homology. For example, the predicted periplasmic *c*-type cytochromes ExtA, ExtG, ExtK, and OmaB vary in size from 25 to 72 kDa, contain 5 to 15 hemes, and share 18%-26% identity (Fig. 1B).

To screen for physiological roles of these loci, single cluster mutants were first constructed in an isogenic background, comprising Δ*extABCD*, Δ*extEFG*, Δ*extHIJKL*, and Δ*ombB-omaB-omcB-orfS-ombC-omaC-omcC* (abbreviated as Δ*omcBC)* mutants. Previous studies have reported complementary roles of OmcB and OmcC (43, 45), thus the entire *omcBC* cluster was removed to screen for conditions under which this pair of homologous conduits were necessary. As these single mutant strains lacked any antibiotic cassettes, they could be used as backgrounds for further double and triple deletions. Multiple cluster deletion mutants leaving only one conduit cluster on the genome are referred to by their remaining cluster, e.g. *“extABCD^+^”* contains only *extABCD* and is constructed by Δ*extEFG* Δ*extHIJKL* Δ*omcBC*, while the mutant lacking all *extABCD, extEFG, extHIJKL, omcB-based* and omcC-based clusters is referred to as “Δ5”. After whole-genome resequencing of all terminal strains containing single clusters and the strain missing all clusters (such as *extABCD* ^+^ and Δ5) to verify no off-target mutations accumulated during the many rounds of insertion and recombination, all of these strains were tested under six different extracellular growth conditions varying in solubility, chemical composition, and redox potential.

**Cells lacking single gene clusters have only partial reduction defects with Fe(III) citrate**. Soluble Fe(III) citrate was the simplest extracellular electron acceptor tested in this study, requiring no attachment to a surface, and requiring no appendages such as a pili or secreted cytochromes for reduction. Under these conditions, no single cluster deletion eliminated the majority of soluble Fe(III) citrate reduction. The most severe defect was observed in the Δ*omcBC* cluster mutant, which grew slower than any other single mutant and reduced only 60% of Fe(III) citrate compared to wild type (Fig. 2A). Minor defects were also observed for Δ*extEFG* and Δ*extHIJKL*, while Δ*extABCD* reduced Fe(III) citrate at wild-type levels. These results suggested that more than one cluster was necessary for wild type soluble Fe(III) reduction.

**Figure 2.**
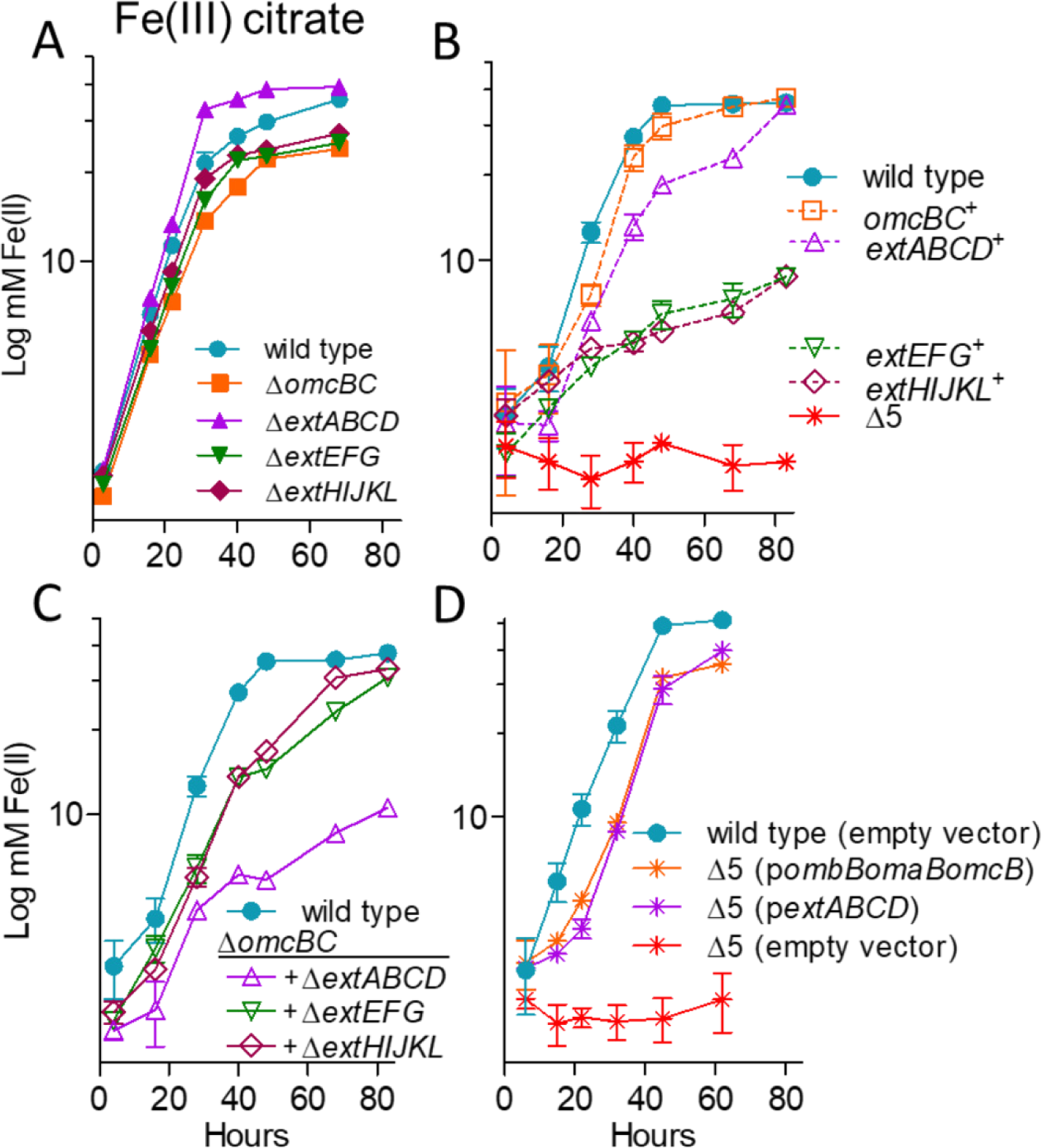
OmcBC or ExtABCD are sufficient during Fe(III)-citrate reduction, deletion of all clusters eliminates Fe(III)-citrate reduction. Growth using 55 mM Fe(III)-citrate as an electron acceptor by A) single conduit cluster deletion mutants, B) triple mutants lacking all but one cytochrome conduit, as well as the Δ5 strain lacking all five cytochrome conduits, C) mutants in an *ΔoμcΒC* background strain, and D) Δ5 mutants expressing *omcB* or *extABCD* or carrying an empty expression vector as control. All experiments were conducted in triplicate and curves are average ± SD of n ≥ 3 replicates.

**Any one gene cluster is sufficient for partial Fe(III) citrate reduction, and deletion of all 5 clusters eliminates electron transfer to this substrate**. Deletion of the full suite of clusters was the only combination that eliminated all residual electron transfer to Fe(III) citrate (Fig. 2B). When multiple-deletion strains still containing one cluster were tested for Fe(III) citrate reduction, results supported key roles for *omcBC* and *extABCD* in soluble metal reduction, and little involvement by *extEFG* or *extHIJKL.* For example, Fe(III) citrate reduction by *omcBC* ^+^ and *extABCD* ^+^ was comparable to that of wild type, while *extEFG* ^+^ and *extHIJKL* ^+^ strains reduced Fe(III) citrate to just 20% of wild type.

**Only strains lacking both *omcBC* and *extABCD* had a significant defect in Fe(III) citrate reduction**. Because Δ*omcBC* demonstrated the largest defect in Fe(III) citrate reduction, additional deletions in this background were tested for their ability to reduce this substrate (Fig. 2C). Only the double cluster deletion mutant Δ*omcBC* Δ*extABCD* reduced Fe(III) citrate at a significantly lower rate compared to the Δ*omcBC* strain, which agreed with the robust growth seen in strains containing only *omcBC* ^+^ or *extABCD* ^+^. The Δ*omcBC* Δ*extABCD* mutant (still containing both *extEFG* and *extHIJKL)* reduced Fe(III) citrate poorly, to the same level as their single-cluster strains containing only *extEFG* ^+^ or *extHIJKL* ^+^ (Fig. 1B vs. Fig. 2C). These data suggested that when both *extEFG* and *extHIJKL* remained in the genome, their contribution was not additive.

Not shown in Fig 2 is metal reduction data for intermediate deletion mutants with no additional phenotype such as Δ*extEFG* Δ*extHIJKL.* Experiments performed after such double mutants were constructed revealed no changes that deviated from wild type or their parent single-cluster deletions. Only intermediate strains with additive phenotypes, such as strains in the Δ*omcBC* background, are shown in Fig 2.

**Expression of single conduit clusters from vectors is sufficient to recover Fe(III) citrate reduction**. When compared to empty-vector controls, complementation of the Δ5 strain with single *omcB* (as *ombB-omaB-omcB*) or *extABCD* clusters restored Fe(III) citrate reduction to levels within 90% of the respective *omcBC* ^+^ and *extABCD* ^+^ strains (Fig. 2D). Previous studies have also shown that expression of only the omcB-based cluster is sufficient to rescue ferric citrate reduction defects in a Δ*omcBC* strain (45), but *extABCD* has never been used to rescue a respiratory phenotype. These data are the first evidence that a putative outer membrane complex other than those encoded in *omcB* could be sufficient for extracellular metal reduction in *Geobacter.*

**Only strains lacking multiple gene clusters have significant defects in Fe(III)- and Mn(IV)-oxide reduction**. Particulate metal oxides represent substrates of additional complexity, requiring pili and additional cytochromes for long-range electron transfer to particles or surfaces after transmembrane electron transfer. Because they are not hypothesized to act as the interface with distant electron acceptors, it was possible the outer membrane complex mutants would show less specificity during reduction of Fe(III) or Mn(IV) oxides. However, trends remained similar to Fe(III) citrate, where deletion of single conduit clusters in *G. sulfurreducens* only had modest effects on metal oxide reduction (Fig. 3A and C) and additional conduit cluster deletions were needed to severely impact growth (Fig. 3B and D). The most severe defect was again observed in the Δ*omcBC* cluster mutant, which reduced 68% of Fe(III)-oxide compared to wild type (Fig. 3A). Minor defects were observed for single Δ*extEFG* and Δ*extHIJKL* deletions, while Δ*extABCD* reduced Fe(III)-oxide near wild-type levels. In contrast, none of the single mutants displayed defects with Mn(IV)-oxides (Fig. 3C).

**Figure 3.**
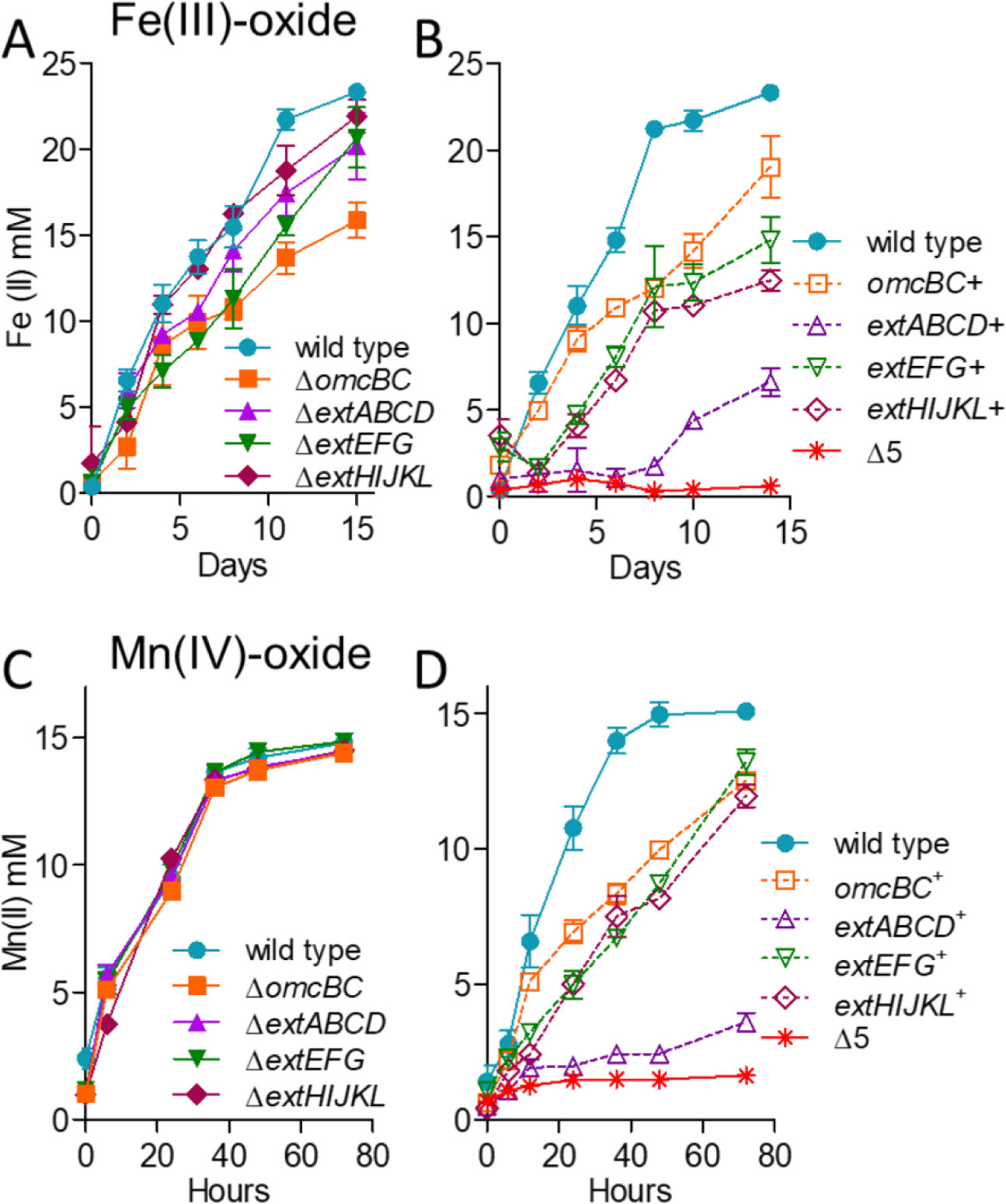
No single outer membrane cluster is essential but all are necessary for wild type levels of electron transfer to Fe(III)- and Mn(IV)-oxides. Growth of single cluster deletion mutants and triple mutants lacking all but one cytochrome conduit cluster, as well as the Δ5 mutant lacking all clusters utilizing A) 70 mM Fe(III)-oxide or B) 20 mM Mn(IV)-oxide as terminal electron acceptor. All experiments were conducted in triplicate and curves are average ± SD of n ≥ 3 replicates.

Unlike soluble metal reduction, however, results supported roles for *omcBC* and *extEFG* in metal oxide reduction and little involvement by *extABCD.* For example, in strains containing only one cluster, Fe(III)-oxide reduction by *omcBC* ^+^ was nearly 80% of wild type, *extEFG* ^+^ was over 60%, but the *extABCD* ^+^ strain reduced less than 30% of wild type. Similarly, the *omcBC* ^+^, *extEFG* ^+^, and *extHIJKL* ^+^ strains achieved about 80% of wild type Mn(IV)-reduction at 80 hours, but the *extABCD* ^+^ strain again displayed poor Mn(IV)-oxide reduction. As with soluble metal reduction, deletion of the full suite of clusters was necessary to eliminate all residual electron transfer to either Fe(III)- or Mn(IV)-oxides (Fig. 3B and D).

**Only strains lacking both *omcBC* and *extEFG* had a significant defect in Fe(III)- and Mn(IV)-oxide reduction**. Since Δ*omcBC* demonstrated the largest defect in Fe(III)-oxide reduction, additional deletions in this background were tested during Fe(III) and Mn(IV)-oxide reduction (Fig. 4). Fe(III)-oxide reduction by Δ*omcBC* Δ*extEFG* was less than 25% of wild type, while the Δ*omcBC* Δ*extABCD*, and Δ*omcBC* Δ*extHIJKL* strains still reduced Fe(III)-oxides similar to the Δ*omcBC* strain. The additive effect from Δ*extEFG* agreed with data from mutants containing single clusters, where *omcBC* ^+^, and *extEFG* ^+^ showed the best reduction. The Δ*omcBC* Δ*extEFG* strain also had a severe Mn(IV)-oxide reduction defect. However, unlike Fe(III)-oxide reduction, the Δ*omcBC* Δ*extABCD* and Δ*omcBC* Δ*extHIJKL* double deletion strains had only a modest Mn(IV) reduction defect, suggesting higher contributions of *extABCD* and *extHIJKL* clusters during Mn(IV) compared to Fe(III) reduction.

**Figure 4.**
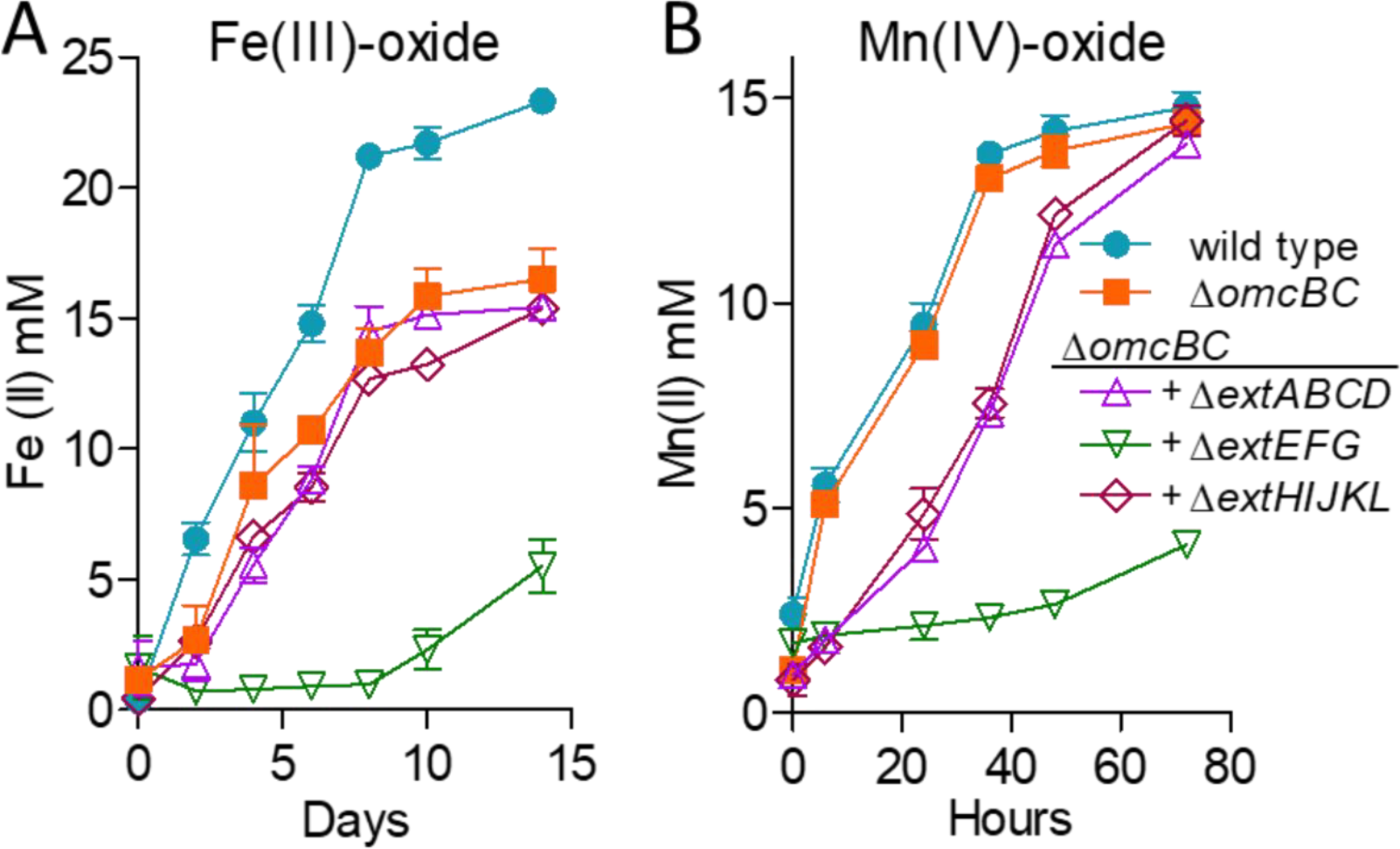
OmcBC and ExtEFG have additive roles in Fe(III)- and Mn(IV)-oxide reduction. Reduction of A) 70 mM Fe(III)-oxide or B) 20 mM Mn(IV)-oxide by the *ΔomcBC* strain and additional deletions in an *ΔomcBC* background. All experiments were conducted in triplicate and curves are average ± SD of n ≥ 3 replicates.

The poor growth of the Δ*omcBC* Δ*extEFG* mutant with insoluble metals was surprising, since this strain still contained *extHIJKL.* When *extHIJKL* was the only cluster remaining, the *extHIJKL*^+^ strain reduced up to 50% of Fe(III)-oxide and 75% of Mn(IV)-oxide compared to wild type (Fig. 3B and D; Table 2). This was a rare case where performance of a mutant containing the single cluster performed better than predicted by single and double mutants, and raises the hypothesis that *extHIJKL* expression or function is partially inhibited by the presence of *extABCD.* No other *ext* or *omc* cluster showed this kind of behavior with soluble or insoluble metals.

**Expression of single conduit clusters partially recovers Fe(III)- and Mn(IV)-oxide reduction**. Plasmids containing constiutive *ombB-omaB-omcB* or *extABCD* clusters resulted in partial recovery (Fig. 5), consistent with the intermediate phenotypes displayed by mutants retaining these single clusters on the genome. Expression of the *omcB* cluster reestablished Fe(III)-oxide reduction to a level less than that seen in the *omcBC* ^+^ strain containing the full duplicated cluster in its original context, suggesting that both *omcB* and *omcC* are necessary (Fig. 4B). Expressing *extABCD* from a plasmid restored Fe(III)-oxide reduction in the Δ5 strain near the low levels of the *extABCD* ^+^ strain, and reduction of Mn(IV)-oxides by *omcB* or extABCD-expressing strains was even lower. These data again agreed with the partial reduction phenotype of mutant strains containing only *extABCD.*

**Figure 5.**
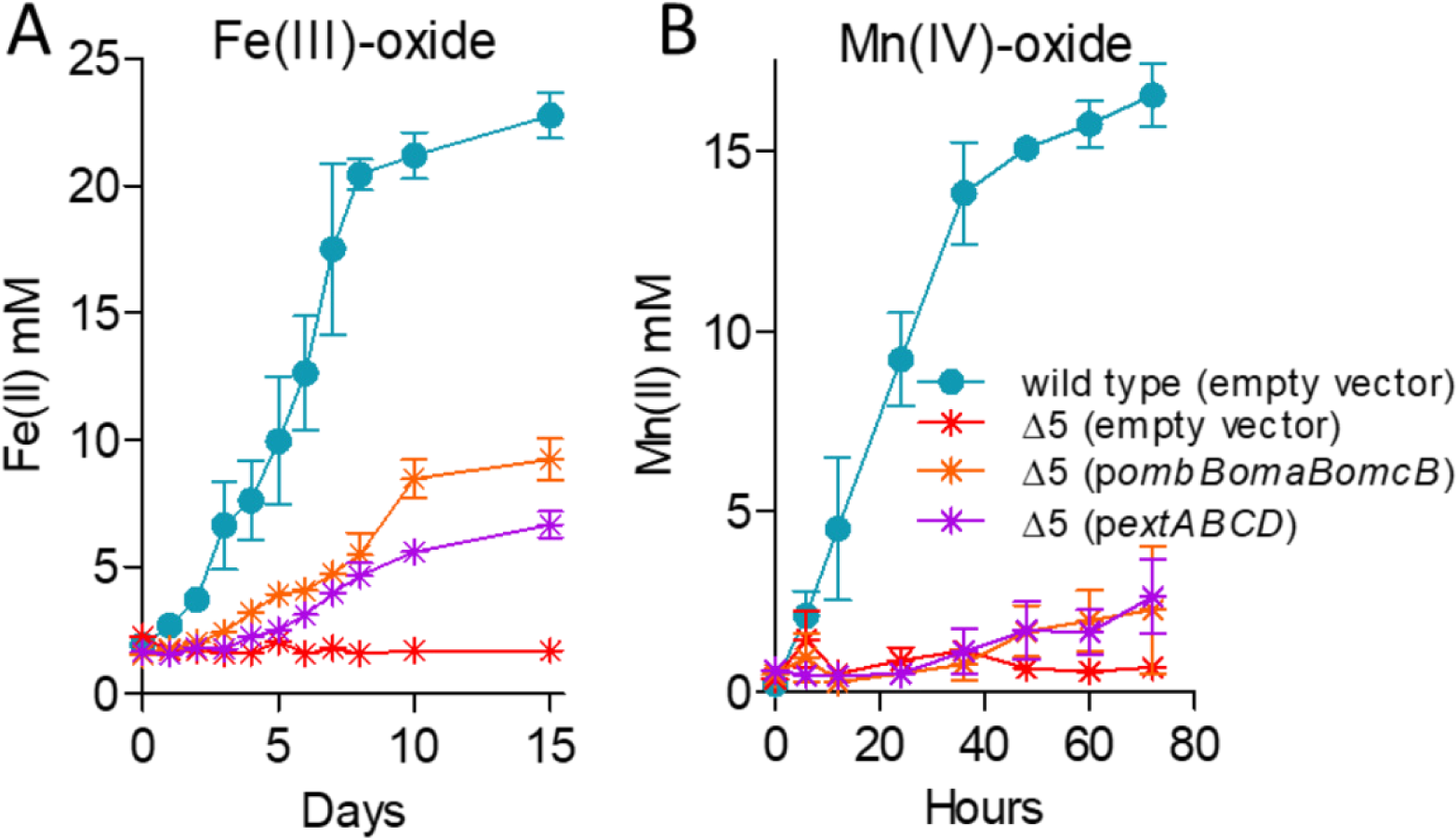
Partial complementation by single conduit clusters supports hypothesis that multiple conduit complexes are necessary for wild-type levels of metal oxide reduction. Reduction of A) 70 mM Fe(III)-oxide or B) 20 mM Mn(IV)-oxide by the Δ5 mutant expressing *extABCD* or the *omcB* cluster compared to the empty vector control. All experiments were conducted in triplicate and curves are average ± SD of n ≥ 3 replicates.

**Mutants lacking *extABCD* are defective in electrode growth at all redox potentials, while mutants containing only *extABCD* outperform wild type**. In contrast to metal reduction, when strains were grown as biofilms on electrodes poised at high (0.24 V vs. SHE) or low (−0.1 V, (46)) redox potentials, only Δ*extABCD* mutants showed a defect in both the rate and extent of growth. Mutants lacking the *omcBC* and *extEFG* clusters grew at similar rates as wild type, while Δ*extHIJKL* demonstrated a lag before growing with a similar doubling time as wild type (Fig. 6A). In all experiments, Δ*extABCD* grew poorly, without a clear exponential phase. The apparent doubling time of Δ*extABCD* was longer than 20 h, or over 3-fold slower than wild type, and only reached 20% of wild type final current density, or 116 ± 33 μA/cm^2^ vs. 557 ± 44 μA/cm^2^ (n ≥ 5 per strain).

**Figure 6.**
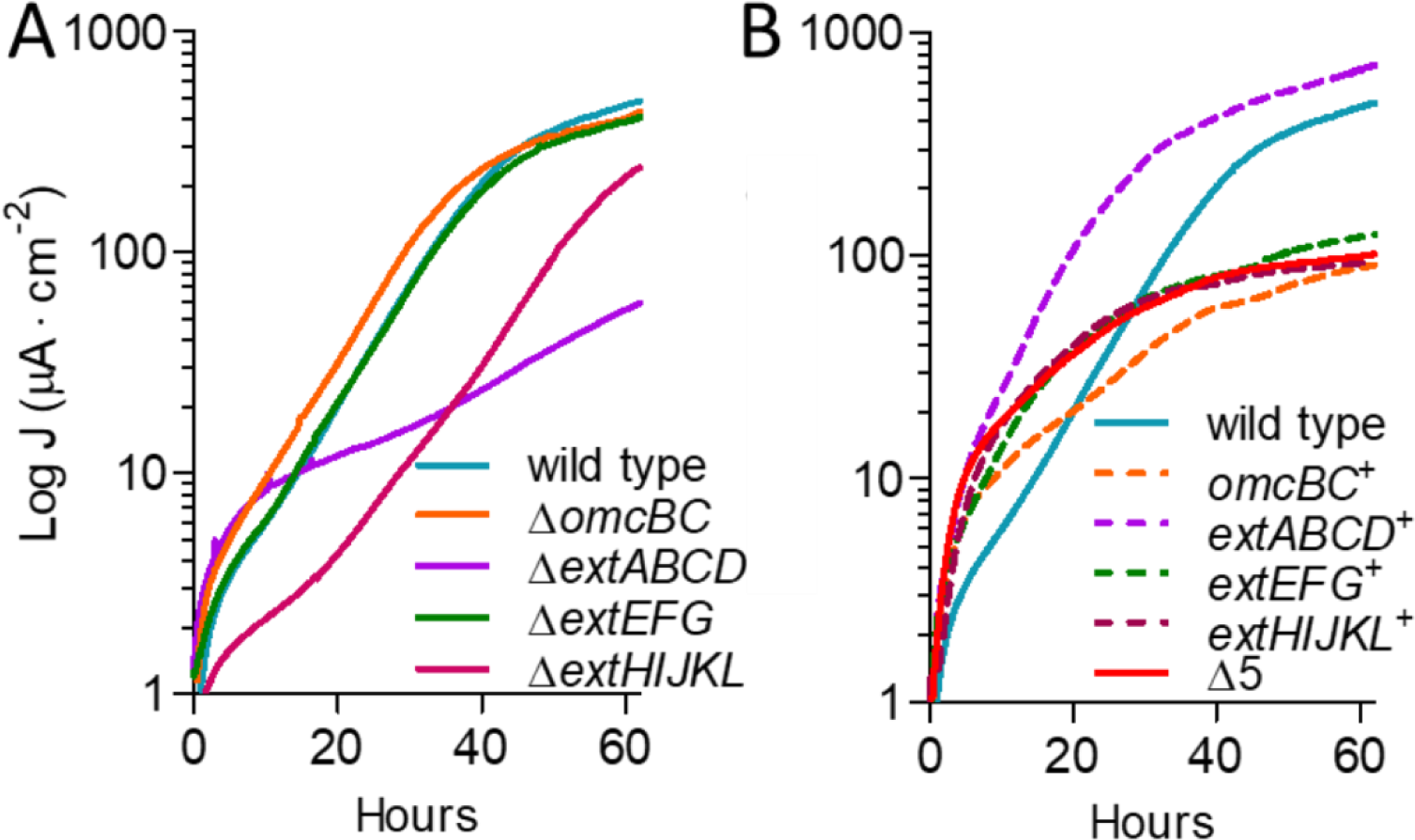
Only the ExtABCD conduit cluster is necessary for electrode reduction. Current density produced by A) single and B) multiple-cluster deletion mutants on graphite electrodes poised at +0.24 V vs. SHE. All mutants were grown in at least two separate experiments, and curves are representative of n ≥ 3 independent replicates per experiment. Similar results were obtained at lower (−0.1 V vs. SHE) redox potentials.

Mutants containing only one gene cluster *(extABCD* ^+^, *extEFG* ^+^, *extHIJKL*^+^, *omcBC* ^+^) as well as a mutant lacking all gene clusters (Δ5) were then analyzed for growth on electrodes. The Δ5 mutant grew at the same low, nonexponential rate as the Δ*extABCD* single mutant at both redox potentials, suggesting that none of the additional clusters were responsible for residual growth rate originally seen in *ΔextABCD.* In contrast, *extABCD* ^+^ grew faster than wild type (4.5 ± 0.2 h vs. 6.5 ± 0.3 h doubling time, n ≥ 9) density 40% higher than wild type (768 ± 52 μΑ/cm^2^ vs. 557 ± 44MA/cm^2^, n≥9). All other multiple-deletion strains containing only one cluster grew as poorly as the Δ5 mutant, further indicating that under these conditions, *extEFG, extHIJKL*, and *omcBC* were not necessary or sufficient to restore electron transfer to electrodes (Fig. 6B). We were unable to identify the origin of the slow growth enabling residual electron transfer to electrodes, although *G. sulfurreducens* contains at least 3 other multiheme cytochrome-rich regions with conduit-like signatures that remain to be examined.

**A 5-conduit deletion mutant expressing *extABCD* has a faster growth rate on electrodes than wild type**. To further investigate the specific effect of *extABCD* on electrode growth, *extABCD* was provided on a vector in the Δ5 strain. The 3-gene *omcB* conduit cluster *(ombB-omaB-omcB)* was also placed in the Δ5 strain using the same vector, and both were compared to wild type cells containing the empty vector. While the plasmid is stable for multiple generations, routine vector maintenance requires growth with kanamycin, and kanamycin carry-over into biofilm electrode experiments is reported to have deleterious effects on electrode growth (23, 56). Thus, we first reexamined growth of the empty vector strain. When selective levels of kanamycin (200 μg·ml^−1^) were present in electrode reactors, colonization slowed and final current production decreased 74% even though cells carried a kanamycin resistance cassette. At levels resulting from carry-over during passage of cells into the electrode reactor (5 μg·ml^−1^) growth rate of vector-containing cells was not affected, but final current was decreased up to 30%, suggesting interference with biofilm formation rather than respiration (Fig. 7A). All subsequent complementation was performed in the presence of 5 μg·ml^−1^ residual kanamycin and compared to these controls.

**Figure 7.**
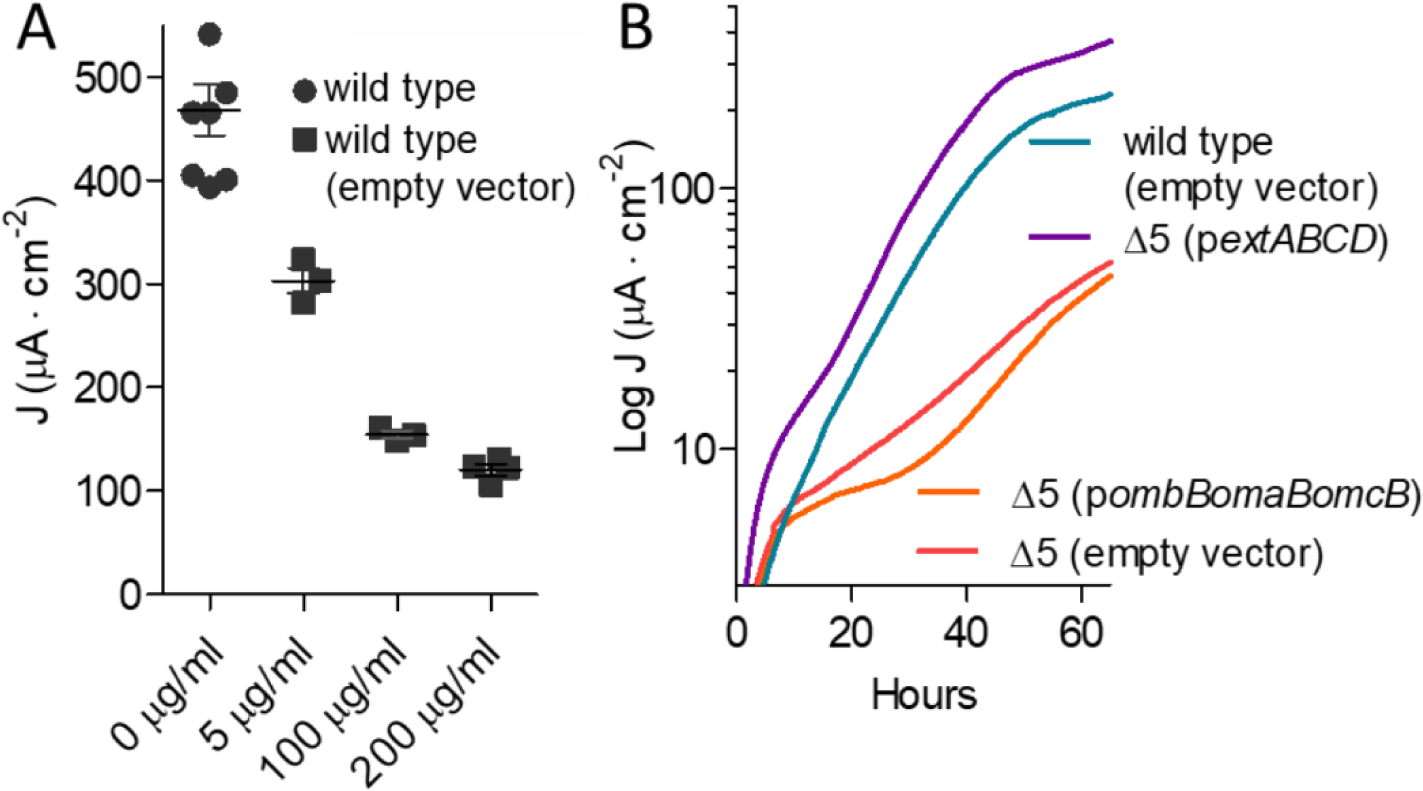
Effect of kanamycin on final current density, and comparison of ExtABCD and OmcBC complementation. A) Final current density of wild type *G. sulfurreducens* compared to wild type carrying an empty vector in the presence of increasing kanamycin concentrations. B) Current density produced by the Δ5 strain plus either *extABCD* or *omcB* cluster-containing vectors, in the presence of 5 μg/ml residual kanamycin. Wild type and Δ5 strains carrying the empty vector were used as controls. All experiments were conducted in duplicate and curves are representative of n ≥ 3 replicates per experiment.

Expressing the *omcB* conduit cluster in the Δ5 strain failed to increase growth with electrodes as electron acceptors. These data were consistent with the lack of an effect seen in *ΔomcBC* deletions, and the poor growth of *omcBC* ^+^ mutants that still contained both the OmcB and OmcC clusters in their native genomic context (Fig. 7B). In contrast, when *extABCD* was expressed on the same vector in the Δ5 background, colonization was faster and cells reached a higher final current density compared to wild type carrying the empty vector (421 ± 89 μA/cm^2^ vs. 297 ± 11 μA/cm^2^, n=3) (Fig. 7B). This enhancement by plasmid-expressed *extABCD* (141% of wild type with empty vector) was similar to the positive effect observed in the *extABCD* ^+^ strain (137% of wild type) (Fig. 6B), and further supported the hypothesis that *extABCD* is both necessary and sufficient during growth with electrodes.

Growth of any two-conduit deletion mutant was unchanged from single-cluster strains (Fig. S1). For example, just as the mutant lacking *extABCD* produced the same phenotype as the Δ5 strain (Fig. 6), deletion of a second cluster from the Δ*extABCD* strain produced similar results as Δ*extABCD* alone, and no two-cluster combination of *omcBC, extEFG* or *extHIJKL* showed defects to suggest they were required during electrode growth conditions, or to indicate their presence affected expression of *extABCD.* The Δ*extABCD* and Δ5 strains were also monitored during extended incubation times to determine if final current density increased after a prolonged incubation period, but current remained unchanged even after 200 h (Fig. S2).

**Transcriptomic analysis reveals no differential expression of putative conduit clusters during growth on electrodes, or off-target expression effects in *extABCD* ^+^.** The importance of the *extABCD* gene cluster during electrode growth was first discovered via genetic experiments (46), but none of the *ext* genes described here were highlighted or examined in earlier studies measuring transcriptional or proteomic changes. Data is available from microarray studies comparing stationary phase electrode biofilms with >4 day old fumarate biofilms grown under electron donor limitation (24), or comparing stationary phase electrode biofilms with Fe(III) citrate grown cells (57). As mature biofilms contain many layers of inactive or slowly growing cells (58), we conducted new experiments capturing both fumarate- and electrode-grown cells during exponential growth to determine absolute levels of transcriptional abundance for *ext* and *omc* genes, using RNAseq.

Figure 8A compares expression levels of wild type *G. sulfurreducens* during exponential fumarate growth to exponential growth with electrodes, using data averaged from at least 2 biological replicates under each condition. Despite the fact that this represents a shift from planktonic cells using an intracellularly reduced acceptor to biofilms using an extracellular acceptor, few genes undergo changes +/− Log_2_ > 2. Highlighted in Fig 8A are all annotated cytochromes and pili genes reported to be involved in metal or electrode respiration, showing that nearly all of these were constitutively expressed between the two laboratory conditions of non-limiting electron acceptor.

**Figure 8.**
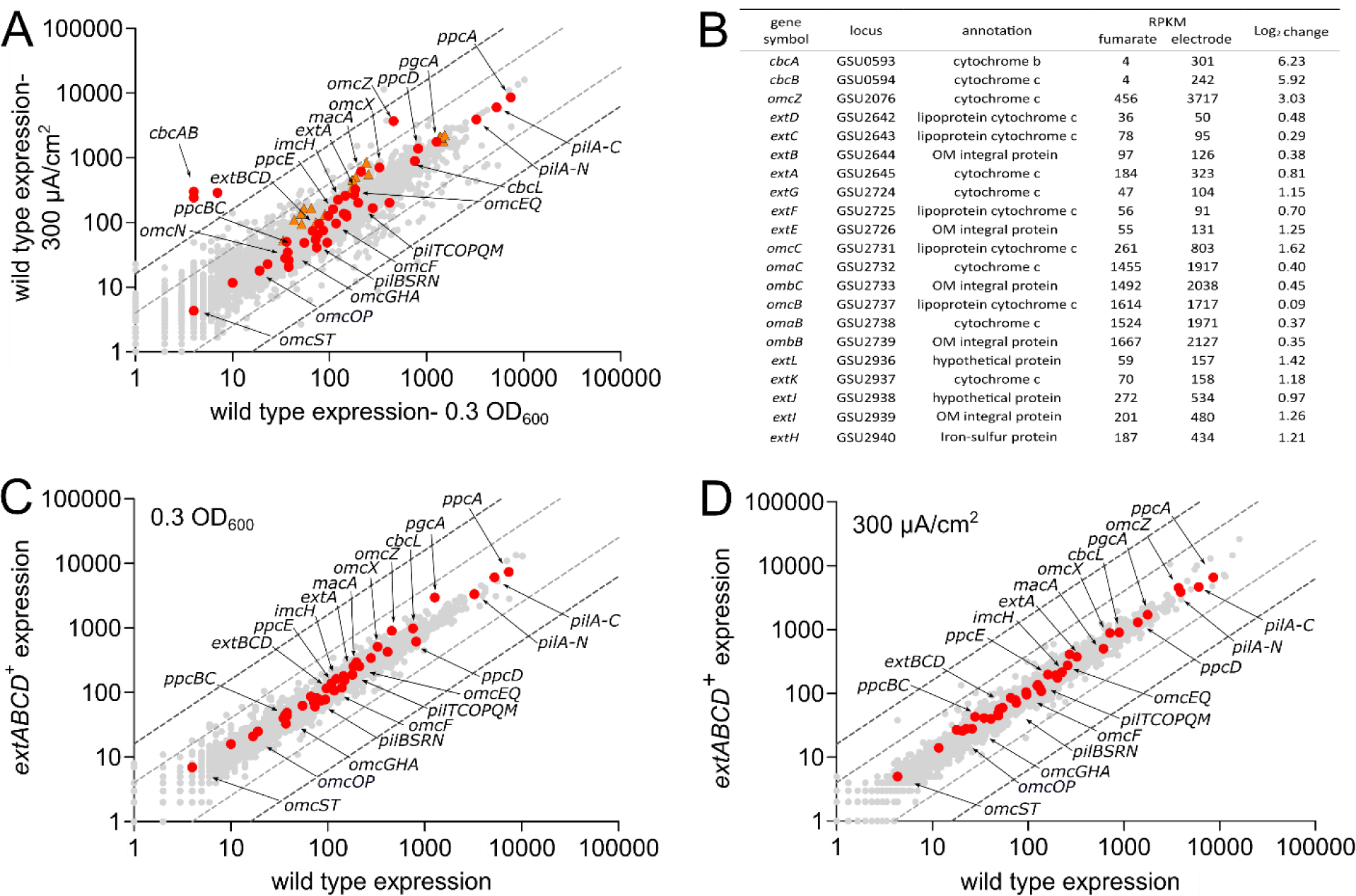
Transcriptomic analysis shows no significant differences in expression between *extABCD*^*+*^ and wild type strains̤. A) Comparison of expression levels of wild type exponentially growing cells under fumarate- and electrode-respiring conditions, showing no significant up- or downregulation of *ext* clusters.(orange triangles) or most other known electron transfer proteins (red circles). Dark and light gray dotted lines represent thresholds of 4 and 2 Log2, respectively. B) RPKM and Log2 change of ORFs with largest expression changes as well as genes studied in this work (for additional data, see Table S2). Comparison of the transcriptome of wild type and *extABCD* ^+^ cells exponentially growing using C) fumarate or D) electrode poised at +240 mV as terminal electron acceptor, showing no changes to electron transfer proteins due to deletion of *omBC, extEFG*, and *extHIJKL* clusters. Averages shown of biological replicate samples.

Compared to the highly expressed *omcBC* genes, genes for *extABCD* and other *ext* clusters were expressed at levels equivalent to only 10-20% of *omcB* under both conditions, which may explain OmcB’s dominance in prior gel-based heme stain identification and proteomic analyses. We did observe an overall trend of increased cytochrome and electron transfer gene expression during growth on electrodes, reflecting a general increase in extracellular respiratory processes, but these changes occurred in both essential and nonessential genes. Genes encoding the characterized inner membrane electron transfer proteins ImcH or CbcL also did not change significantly in expression between these two conditions, nor did any genes for periplasmic cytochromes or pili components known to be essential (18, 22, 23). As has been shown before (24), the well-characterized extracellular cytochrome *omcZ* was upregulated over 8-fold during electrode reduction, and a putative inner membrane *c*- and *b*-type cytochrome similar to CbcL that is up-regulated during Fe(III) reduction *(cbcBA)* also increased over 30-fold (51) (Fig. 8A). A table providing RPKM data for all genes studied in this paper, along with *omcZ* and *cbcBA*, is provided in Fig 8B. As none of the *extABCDEFGHIJKL* genes were strongly induced or repressed during electrode growth, and these genes were generally expressed at levels 1/10^th^ of the *omcBC* locus, their absence from prior differential expression analyses is understandable.

A second question that often arises in the study of complex phenotypes is whether deletion of an important or highly expressed cluster such as *omcBC* affects expression of other genes, especially as phenotypes such as biofilm growth require secretion of complexes to the outer membrane, adhesion of cells to surfaces, and production of extracellular proteins such as pili (59–61). The fact that the *extABCD* ^+^ strain lacking 15 different genes always grew faster than wild type, and produced more current than wild type, raised a significant question regarding possible off-target effects on other aspects of metabolism. Therefore, the transcriptome of the *extABCD* ^+^ strain was analyzed under both fumarate- and electrode-respiring conditions, and compared to wild type.

No significant increase or decrease in expression of any previously studied electron transfer proteins were found during growth in fumarate, or during exponential growth on electrodes, when the *extABCD* ^+^strain was compared to wild type (Fig. 8C-D). This further suggested the increased growth rate was not due to higher expression of an unknown gene enabling electron transfer or attachment. It also underscored the trend in *Geobacter* that many genes such as *omcB* are among the most highly expressed under laboratory conditions, yet these expression levels have not correlated with essentiality or function. The full data sets plotted on Figure 8 can be found in Table S2.

**Summary of phenotypes for all Omc and Ext electron conduit gene cluster mutants**. Table 2 summarizes all extracellular reduction phenotypes of single cluster deletions and deletions leaving only one conduit on the genome, adjusted to wild type performance. Each gene cluster was necessary under different conditions. Many of the recently described *ext* gene clusters were necessary for wild-type metal reduction, yet few were sufficient. For example, *extEFG* and *extHIJKL* were necessary for Fe(III) citrate reduction, as strains lacking these clusters only reduced ~65% of wild type levels. But when only *extEFG* or only *extHIJKL* was present, they were not sufficient to reduce Fe(III) citrate at wild type levels. In contrast, the *omcBC* cluster or the *extABCD* cluster alone was necessary for Fe(III) citrate reduction, and the *extABCD* cluster alone was also sufficient for electrode growth. Deletion of all five conduit clusters resulted in complete elimination of metal reduction abilities, while some residual activity remained when the same Δ5 strain was grown using electrodes as terminal electron acceptor. These comparisons show each gene cluster is necessary under at least one of the conditions studied, and provides evidence for additional undiscovered mechanisms enabling transmembrane electron transfer during electrode growth.

## DISCUSSION

Sequencing of the *G. sulfurreducens* genome revealed an unprecedented number of electron transfer proteins, with twice as many genes dedicated to respiratory and redox reactions as organisms with similarly-sized genomes (62). Out of 111 *c*-type cytochromes, 43 had no known homolog, and many were predicted to reside in the outer membrane. The large complement of outer membrane redox proteins in *G. sulfurreducens* became even more of an anomaly as the simpler electron transfer strategy of metal-reducing *S. oneidensis* emerged. If *Shewanella* only requires a single inner membrane cytochrome and a single outer membrane conduit to reduce a multitude of substrates (36, 39, 40, 53), why does *Geobacter* have so many cytochromes?

Evidence that more than one *G. sulfurreducens* outer membrane pathway exists for reduction of extracellular substrates has accumulated in at least 11 separate studies since discovery of OmcB (34, 43, 45). Deletion of *omcB* impacted Fe(III)-reduction, but had little effect on U(IV) or Mn(IV)-oxide reduction (51, 63). A Δ*omcB* suppressor strain evolved for improved Fe(III)-citrate growth still reduced Fe(III)-oxides poorly (44). Strains lacking *omcB* grew similar to wild type on electrodes in four different studies, (24, 29, 57, 64), and OmcB abundance was shown to be lowest on cells near electrodes (65). An insertional mutant lacking six secreted or outer membrane-associated cytochromes in addition to OmcB still demonstrated Fe(III)-oxide reduction (66). After replacing the entire *omcBC* region with an antibiotic cassette and still finding residual Fe(III)-reducing ability, Liu *et al.* (2015) speculated that other *c*-type cytochrome conduit-like clusters in the genome might be active. Most recently, Tn-seq analysis of electrode-grown cells found little effect of *omcB* cluster mutations, yet noted significant defects from insertions in unstudied clusters with *c*-type cytochrome features (46). This combined evidence led us to seek alternative conduit gene clusters that could address both the longstanding mystery of growth by *omcB* mutants and the complexity of electron transfer proteins in the *Geobacter* genome.

The genetic analysis presented here supports a role for these unstudied conduit gene clusters during extracellular respiration. All mutants still containing at least one cluster retained a partial ability to reduce metals, while deletion of the entire *omcBC* region, plus all three *ext* clusters, finally was able to eliminate metal reduction. This need to delete more than one cluster helps explain variability reported with other mutants, and the rapid evolution of suppressors in Δ*omcB* mutants.

In the case of electrodes at both high and low potential, only deletion of *extABCD* altered phenotypes. Additionally, a strain with only *extABCD* remaining on the genome outperformed wild type in terms of growth rate and final current density when grown on electrodes. Since expression of *extABCD* was also able to restore reduction of the soluble acceptor Fe(III) citrate, this cluster can confer the phenotype of extracellular respiration under a condition where pili and secreted cytochromes are not known to be important. Overall, these data show that for all tested metal acceptors, more than one conduit cluster is necessary for wild type levels of reduction, any one cluster can support partial reduction of may metals, and only one cluster can be linked to electrode respiration.

Genetic analyses are typically a first step, designed to reveal which genes are necessary for a phenotype, and worthy of further study. Biochemical and biophysical analyses will be needed to prove (1) if products of *ext* gene clusters indeed function as conduits to transfer electrons across the outer membrane, and (2) identify the proteins or metals these complexes interact with to explain why these clusters seem so tightly linked to growth with certain substrates. Expression analyses failed to detect large differences in *ext* or *omcBC* family genes during transitions between acceptors, arguing against changes in expression as an explanation for specificity. Our ability to complement growth with electrodes in the Δ5 mutant by expressing *extABCD* from a vector, while the *omcB* conduit could not complement growth, further argues against expression differences causing these phenotypes. Unknown post-transcriptional events could be caused by the absence of different gene clusters, but the genetic conclusion that these gene clusters are necessary remains the same.

To reduce metal particles or surfaces likely requires each membrane-bound complex to interact with extracellular proteins such as OmcZ, OmcS, PgcA, or pili, to aid transfer of electrons to the final destination. If these partner proteins are not expressed or made available under all conditions, an outer membrane complex may not be capable of contributing to respiration. In the case of soluble metals such as Fe(III) citrate, conduit complexes should be able to directly reduce the acceptor, making apparent specificity more likely due the ability of the complex(es) to interact with Fe(III) directly.

It is also important to consider lessons from insertional deletions in *G. sulfurreducens*, such as the diheme peroxidase MacA. Initially hypothesized to be an inner membrane quinone oxioreductase, based on the defective phenotype of Δ*macA* mutants during Fe(III)-citrate reduction (67), this phenotype was later explained by Δ*macA* mutants not expressing *omcB*, as the Δ*macA* phenotype could be rescued by expressing *omcB* from a vector (68, 69). As MacA is now known to instead be a soluble peroxidase, oxidative stress in early Δ*macA* studied could have resulted in global downregulation of cytochromes. In our work, the availability of every combination of gene cluster deletion and acceptor condition allows many general downregulation hypotheses to be eliminated. For example, if deletion of *extABCD* suppressed production of pili or cytochromes such as OmcS, all Δ*extABCD* mutants would be predicted to show both an electrode and metal oxide defect, which we did not observe.

Initial transcriptomic surveys also failed to find severe or off-target transcriptional effects on known electron transfer proteins from deletion of *ombB-omaB-omcB-orfS-ombC-* omaC-omcC, *extEFG*, or *extHIJKL*, that could explain the enhanced growth of *extABCD* ^+^. The fact that only the *ombB-omaB-omcB* cluster was necessary to restore Fe(III) citrate reduction further indicated that *orfS* was not essential. However, all of these deletions removed many parts of the genome which were not tested for complementation by single genes, leaving open the possibility of regulatory interactions. Also, in a complex system such as this, post-translational events such as polymerization of pilin monomers into filaments, extracellular cytochrome secretion could be affected by the absence of specific proteins under specific conditions. It is difficult to detect negative interactions via RNAseq or proteomic analyses when mutants fail to grow, but such effects should be addressed in future suppressor and heterologous expression studies, now that these clusters have been identified.

**Insights from similar gene clusters in related organisms**. It remains difficult to predict any function for multiheme cytochromes based on sequence alone, so their genetic context may reveal other clues to their role, and aid identification of such clusters in other genomes. None of the *ext* regions fits the pattern of the *mtr* 3-gene ‘cytochrome conduit’ operon of one small (~40 kDa) periplasmic cytochrome, an integral outer membrane protein, and one large (>90 kDa) lipoprotein cytochrome. For example, *extABCD* includes two small lipoprotein cytochromes, *extEFG* is part of a hydrogenase-family transcriptional unit, and *extHIJKL* contains a rhodanese-like lipoprotein instead of an extracellular cytochrome (Fig. 1).

Specifically, the transcriptional unit beginning with *extEFG* includes a homolog of YedY-family periplasmic protein repair systems described in *E. coli* (70), followed by a NiFe hydrogenase similar to bidirectional Hox hydrogenases used to recycle reducing equivalents in Cyanobacteria (71–73). Rhodanase-like proteins related to ExtH typically are involved in sulfur metabolism (74–76) and an outer surface ExtH/rhodanese-like protein is linked to extracellular oxidation of metal sulfides by *Acidithiobacillus ferrooxidans* (77). Deletion of *extI* in *G. sulfurreducens* causes a severe defect in selenite and tellurite reduction (78). These links to metabolism of hydrogen, sulfur, and other oxyanions suggest roles outside of metal reduction, and future genomic searches for electron conduit clusters should consider the possibility of non-cytochrome components, such as FeS clusters, as the exposed lipoprotein.

Now that genes from *ext* operons can be used in searches of other genomes, an interesting pattern emerges in putative conduit regions throughout Desulfuromonadales strains isolated from freshwater, saline, subsurface, and fuel cell environments (Fig. 9). In about 1/3 of cases, an entire cluster is conserved intact, such as *extABCD* in *G. anodireducens, G. soli*, and *G. pickeringii* (Fig. 9B). However, when differences exist, they are typically non-orthologous replacements of the outer surface lipoprotein, such as where *extABC* is followed by a new cytochrome in *G. metallireducens, Geoalkalibacter ferrihydriticus*, and *Desulfuromonas soudanensis.* Conservation of the periplasmic cytochrome with replacement of the outer surface redox lipoprotein also occurs frequently in the *omcB* and *extHIJKL* clusters (Fig 9A and D). For example, of 18 *extHIJKL* regions, 10 contain a different extracellular rhodanese-like protein with *extIJKL*, each with less than 40% identity to *extH.* This remarkable variability in extracellular components, compared to conservation of periplasmic redox proteins, suggests much higher rates of gene transfer and replacement of domains that are exposed to electron acceptors and the external environment.

**Figure 9.**
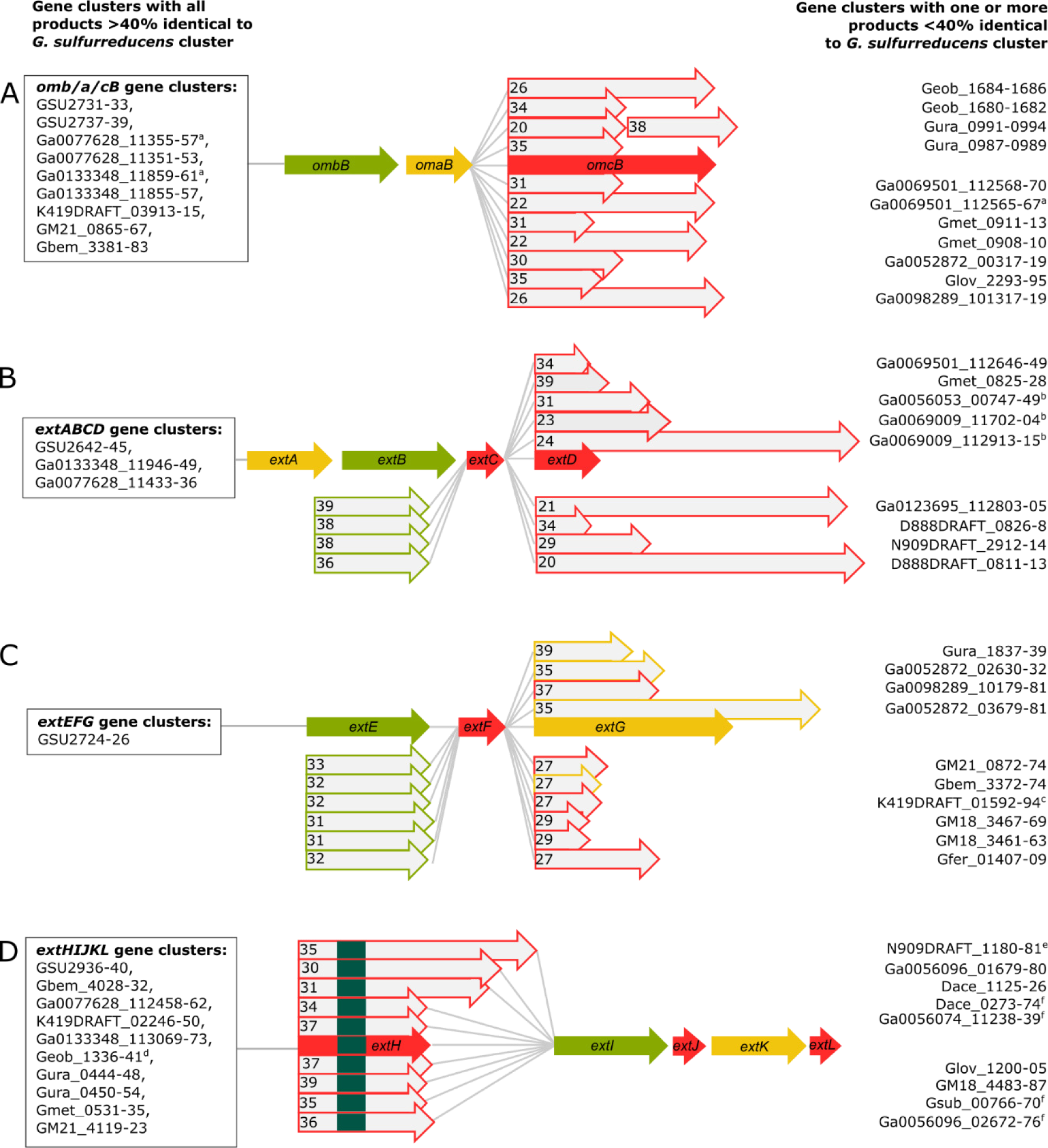
Cytochrome conduit conservation across the Order Desulfuromonodales. Representation of cytochrome conduit clusters from the Desulfuromonodales with homologs to either A) OmcBC, B) ExtABCD, C) ExtEFG, or D) ExtHIJKL. Red arrows = putative outer membrane products with a predicted lipid attachment site, yellow arrows = predicted periplasmic components, green arrows = predicted outer membrane anchor components. Complete clusters with all components sharing >40% identity to the corresponding *G. sulfurreducens* cytochrome conduit are indicated in boxes to the left of each gene cluster. Clusters in which one or more proteins are replaced by a new element with <40% identity are listed on the right side of each gene cluster. Proteins with numbers indicate the % identity to the *G. sulfurreducens* version. ^a^OmcBC homologs in these gene clusters also encode Hox hydrogenase complexes. ^b^Gene clusters have contiguous *extBCD* loci but *extA* is not in vicinity, as *extA* was found in separate parts of the genome for some of those organisms (see Supplemental Table S2). ^c^Gene cluster has additional lipoprotein decaheme *c*-type cytochrome upstream of *extE.* ^d^Lipid attachment sites corresponding to ExtJL could not be found but there is an additional small lipoprotein encoded within the gene cluster. For ExtHIJKL clusters, homologs depicted above *extH* are found in gene clusters containing only *extl*, whereas homologs depicted below *extH* are found in gene clusters containing full *extHIJKL* loci. Upstream and on the opposite strand to all gene clusters homologous to *extHIJKL* there is a transcription regulator of the LysR family, except ^e^, where there is no transcriptional regulator in that region, and ^f^, where there are transcriptional regulators of the TetR family instead.

**Summary**. The data presented here significantly expands the number of genes encoding outer membrane redox proteins necessary during electron transfer in *G. sulfurreducens* and highlights a key difference in the *Geobacter* electron transfer strategy compared to other model organisms. In general, the pattern of multiple genes encoding seemingly overlapping or redundant roles is less like solitary respiratory reductases, and more reminiscent of systems in cellulolytic bacteria that produce numerous similar β-glucosidases to attack a constantly changing polysaccharide substrate (36, 40, 79). A need for multiple outer membrane strategies could be a response to the complexity of metal oxides during reduction; minerals rapidly diversify to become multiphase assemblages of more crystalline phases, the cell:metal interface can become enriched in Fe(II), and organic materials can bind to alter the surface (80–82). Constitutively expressing an array of electron transfer pathways could make cells competitive at all stages with all electron acceptors, allowing *Geobacter* to outgrow more specialized organisms during rapid perturbations in the environment.

## EXPERIMENTAL PROCEDURES

### Growth conditions

All experiments were performed with our laboratory strain of *Geobacter sulfurreducens* PCA as freshly streaked single colonies from freezer stocks. Anaerobic NB medium (0.38 g/L KCl, 0.2 g/L NH4Cl 0.069 g/L NaH2PO4H2O, 0.04 g/L CaCl_2_2H_2_O, 0.2 g/L MgSO_4_7H_2_O, 1% v/v trace mineral mix, pH 6.8, buffered with 2 g/L NaHCO_3_ and flushed with 20:80 N_2_:CO_2_ gas mix) with 20 mM acetate as electron donor, 40 mM fumarate as electron acceptor was used to grow liquid cultures from colony picks. For metal reduction assays, 20 mM acetate was added with either 55 mM Fe(III) citrate, ~20 mM birnessite (Mn(IV)-oxide), or ~70 mM Fe(III)-oxide freshly precipitated from FeCl_2_ by addition of NaOH and incubation at pH 7 for 1 h before washing in DI water. Fe(III)-oxide medium contained an increased concentration of 0.6 g/L NaH_2_PO_4_H_2_O to prevent further crystallization of the metal after autoclaving. All experiments were carried out at 30°C.

### Deletion and complementation construction

Putative conduits were identified through a genomic search for gene clusters containing loci predicted to encode a β-barrel using PRED-TMBB (49), contiguous to periplasmic and extracellular multiheme *c*-type cytochromes or other redox proteins. Localization was predicted by comparing PSORT (48) and the presence/absence of lipid attachment sites (50). Constructs to delete each gene cluster were designed to recombine to leave the site marker-free and also non-polar when located in larger transcriptional units, with most primers and plasmids for the single deletions described in Chan *et al.*, 2017. When genes were part of a larger transcriptional unit or contained an upstream promoter, it was left intact. For example, in the case of the *omcBC* cluster the transcriptional regulator *orfR* (GSU2741) was left intact, and in *extEFG* the promoter and untranslated region was left intact so as to not disrupt the downstream loci.

For deletion mutant construction, the suicide vector pK18*mobsacB* (83) with ~750 bp flanking to the target region was used to induce homologous recombination as previously described (56). Briefly, two rounds of homologous recombination were selected for. The first selection used kanamycin resistance to select for mutants with the plasmid inserted into either up or downstream regions, and the second selection used sucrose sensitivity to select for mutants that recombine the plasmid out of the chromosome, resulting in either wild type or complete deletion mutants. Deletion mutants were identified using a kanamycin sensitivity test and verified by PCR amplification targeting the region. Multiple PCR amplifications with primers in different regions were used to confirm full deletion of each gene cluster (55 and Table S1).

During this work, we found that manipulations in the *omcBC* cluster, which harbors large regions of 100% identity, frequently underwent recombination into unexpected hybrid mutants which could escape routine PCR verification. For example, when *omaB* and *omaC* genes recombined, a large hybrid operon containing *omaB* linked to *ombC-omcC* would result, and sometimes the region would recombine to produce a hybrid of te two repressors controlling expression of the region. Routine primer screening, especially targeting flanking regions, failed to detect the large product. Only via use of multiple internal primers (55 and Table S1), paired with longer-read or single molecule (PacBio) sequencing, were we able to verify and isolate strains in which complete loss of the *omcBC* cluster occurred, and dispose of hybrid mutants. Whole-genome resequencing was also performed on strains containing only one cluster, such as the strain containing only *extABCD*, especially since this strain has an unexpected phenotype where it produced more current than wild type. Because these hybrid *omcBC* operon strains still contained mixed conduits and had altered expression due to disruption of the repressors upstream, verification by PCR and whole genome sequencing (especially with single-molecule techniques able to span the entire ~10 kb region) are recommended to confirm deletions of large and repetitive regions such as the *omcBC* cluster when working with this region.

Mutants lacking a single gene region were used as parent strains to build additional mutations. In this manner, six double gene-cluster deletion mutants, four triple-cluster deletion mutants and one quintuple-cluster deletion mutant lacking up to nineteen genes were constructed (Fig. 1; Table 1). For complementation strains, putative conduits were amplified using primers listed in Table S1 and inserted into the *G. sulfurreducens* expression vector pRK2-Geo2 (56), which contains a constitutive promoter P_*acpP*_. The putative conduit *extABCD* was assembled into a single transcriptional unit to ensure expression.

**Table 1.**
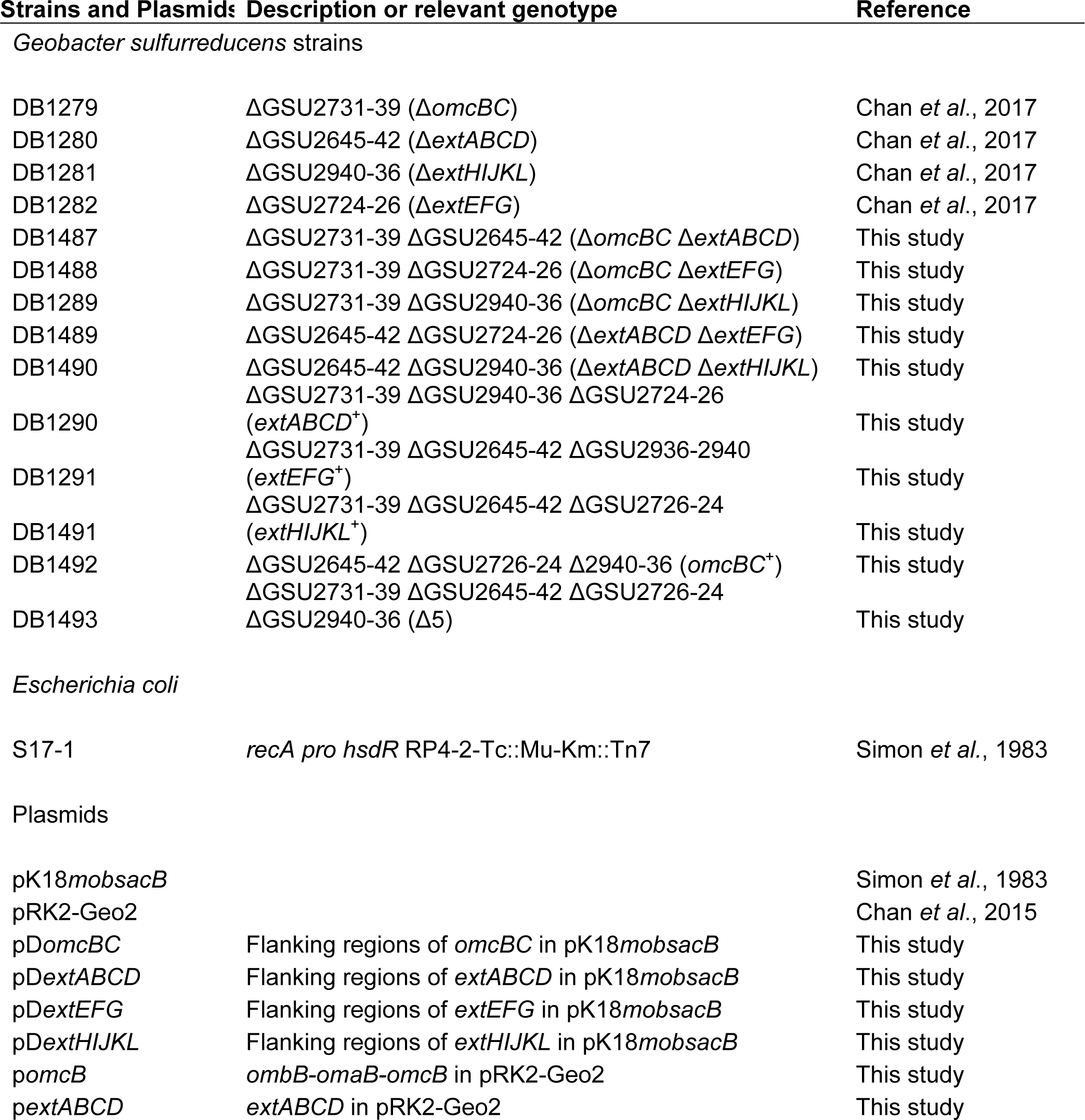
Strains and plasmids used in this study.

**Table 2.** Comparative performance of *G. sulfurreducens* strains lacking one cluster, or containing only one cluster. Growth of single cytochrome conduit deletion mutants and mutants lacking all clusters except one, averaged from eight biological replicates or more and represented as the percent of wild type growth. Averages and standard deviation represented.

| Substrate | % of wild type |  |  |  |  |  |  |  |  |
| --- | --- | --- | --- | --- | --- | --- | --- | --- | --- |
| | $\Delta omcBC$ | $\Delta extABCD$ | $\Delta extEFG$ | $\Delta extHIJKL$ | $omcBC^+$ | $extABCD^+$ | $extEFG^+$ | $extHIJKL^+$ | $\Delta 5$ |
| Fe(III) citrate | 61.2 ± 10.5 | 105 ± 6.6 | 62.5 ± 4.9 | 66.3 ± 2.5 | 101.1 ± 8.4 | 99.2 ± 11.3 | 22.5 ± 2.4 | 23.8 ± 6.4 | 0.1 ± 0.6 |
| Fe(III)-oxide | 68.9 ± 8.4 | 83.3 ± 12.1 | 87.5 ± 14.9 | 95.8 ± 24.9 | 78.8 ± 3.9 | 29.2 ± 2.6 | 60.4 ± 9.5 | 52.1 ± 3.7 | 0.1 ± 0.3 |
| Mn(IV)-oxide | 94.5 ± 6.4 | 95.1 ± 2.8 | 99.6 ± 3.4 | 97.9 ± 6.1 | 83.3 ± 14.1 | 26.7 ± 5.9 | 86.8 ± 6.5 | 75.6 ± 7.3 | 1.7 ± 0.9 |
| Electrode | 76.5 ± 16.5 | 20.9 ± 6.0 | 104.8 ± 2.1 | 86.3 ± 15.3 | 28.3 ± 5.2 | 137.9 ± 9.5 | 21.2 ± 6.5 | 25.9 ± 4.2 | 21.9 ± 4.4 |

### Electrode reduction assays

Sterile three-electrode conical reactors containing 15 mL of NB with 40 mM acetate as electron donor and 50 mM NaCl to equilibrate salt concentration were flushed with a mix of N_2_-CO_2_ gas (80:20, v/v) until the O_2_ concentration reached less than 2 ppm. Liquid cultures were prepared by inoculating 1 ml liquid cultures from single colonies inside an anaerobic chamber. Once these cultures reached late exponential to stationary phase, they were used to inoculate 10 ml cultures with 10% v/v. Each reactor was then inoculated with 25% v/v from this liquid culture as it approached acceptor limitation, at an OD_600_ between 0.48 and 0.52. Working electrodes were set at either −0.1 V or +0.24 V vs SHE and average current density recorded every 12 seconds. Each liquid culture propagated from an individual colony pick served no more than two reactors, and at least three separate colonies were picked for all electrode reduction experiments for a final n ≥ 3.

### Metal reduction assays

NB medium with 20 mM acetate as electron donor and either 55 mM Fe(III)-citrate, ~70 mM Fe(III) oxide, or ~20 mM birnessite (Mn(IV)O_2_) as electron acceptor was inoculated with a 0.1% inoculum of early stationary phase fumarate limited cultures. Time points were taken as necessary with anaerobic and sterile needles. These were diluted 1:10 into 0.5 N HCl for the Fe(III) samples and into 2 N HCl, 4 mM FeSO_4_ for Mn(IV) samples. Samples were diluted once more by 1:10 in the case of Fe(III) assays and by 1:5 in the case of Mn(IV) assays into 0.5 N HCl. FerroZine^R^ reagent was then used to determine the Fe(II) concentration in each sample. Original Fe(II) concentrations were calculated for Fe(III) reduction assays by accounting for dilutions and original Mn(IV) concentrations were calculated by accounting for the concentration of Fe(II) oxidized by Mn(IV) based on the following: Mn(IV) + 2Fe(II) = Mn(II) + 2Fe(III). In other words, two molecules of Fe(II) are reduced by one molecule of Mn(IV). Therefore, the increase of Fe(II) concentration over time in our samples indicates a decrease of Mn(IV), or increase of Mn(II), in a 2:1 ratio.

### RNAseq

For liquid-grown cultures, total RNA was extracted from 10 ml of *G. sulfurreducens* culture grown to mid-log (0.25 - 0.3 OD_600_). For biofilm-grown cultures, total RNA was extracted from graphite electrodes of *G. sulfurreducens* biofilms grown to mid-log (300 μΑ/cm^2^). Biofilms were rinsed to remove planktonic cells and removed from electrodes using a plastic spatula. Cell pellets from all samples were washed in RNAprotect (Qiagen) and frozen at −80°C before RNA extraction using RNeasy with on column DNase treatment (Qiagen). Ribosomal RNA was depleted using RiboZero (Illumina) by the University of Minnesota Genomics Center before stranded synthesis and sequenced on Illumina HiSeq 2500, 125 bp pair-ended mode. Residual ribosomal RNA sequences were removed before analysis using Rockhopper (84). Duplicate biological samples were analyzed for each strain. An in-house re-sequenced *G. sulfurreducens* genome and annotation released in a prior publication was used as reference (46, 56). Full RPKM values are in Table S2, and raw RNAseq reads are deposited in the NCBI SRA under our BioProject PRJNA290373.

### Homolog search and alignment

Homologs to each of the individual cytochrome conduit proteins were queried on 11 −302016 in the Integrated Microbial Genomes database (85) with a cutoff on 75% sequence length and 40% identity based on amino acid sequence within the Desulfuromonadales. A higher percent identity was demanded in this search due to the high heme binding site density with the invariable CXXCH sequence. Only ExtJ and ExtL were excluded from the search and the OmcBC region was collapsed into a single cluster due to the high identity shared between the two copies. The gene neighborhood around each homolog hit was analyzed. With a few exceptions (see Table S2), all homologs were found to be conserved in gene clusters predicted to encode cytochrome conduits and containing several additional homologs to each corresponding *G. sulfurreducens* conduit. The proteins within each homologous cytochrome conduit that did not fall within the set cutoff were aligned to the amino acid sequence of the *G. sulfurreducens* component they replaced using ClustalΩ (86).

## ACKNOWLEDGEMENTS

We thank J. Badalamenti for sequence analysis assistance and data management. This research was supported by the Office of Naval Research (N000141210308). FJO is supported by the National Council of Science and Technology of Mexico (CONACYT).

